# Gain of chromosome region 1q31.3 in human iPSCs confers growth advantage and alters contraction in derivative cardiomyocytes

**DOI:** 10.1101/2023.12.11.571144

**Authors:** Karina O. Brandão, Viviana Meraviglia, Daniela Salvatori, Xu Cao, Luca Sala, Loukia Yiangou, Mervyn P.H. Mol, Milena Bellin, Christine L. Mummery, Richard P. Davis

## Abstract

hPSCs can acquire chromosomal aberrations such as copy number variations during prolonged maintenance *in vitro*, conferring growth advantages. However, the effect of these culture-acquired mutations on the phenotypes of the differentiated hPSCs is largely unstudied. Here, we identified mosaicism in a hPSC line in which some cells showed a gain of chromosome 1q31.3. We subcloned the wild-type and variant hPSCs and could maintain both as stable lines. While both variant and wildtype lines differentiated efficiently to cardiomyocytes (hPSC-CMs), molecular analysis revealed the variant hPSC-CMs had increased expression of *TNNT2*, a gene encoding one of the major sarcomere proteins mediating cardiomyocyte contractility and located within the gained chromosome 1q region. Moreover, the variant hPSC-CMs showed altered contraction kinetics. Together these results highlight the importance of careful monitoring of chromosome aberrations in hPSC lines as these could have confounding effects in various applications such as disease modelling and drug discovery.

## INTRODUCTION

Human pluripotent stem cells (hPSCs) represent a promising avenue for developing human disease models, advancing drug discovery, and serving as potential cell replacement therapies in regenerative medicine. Nonetheless, both human embryonic stem cells (hESCs) and induced pluripotent stem cells (hiPSCs) can acquire genetic changes, such as chromosomal aberrations, copy number variations (CNVs) and point mutations, over extended periods of *in vitro* maintenance and expansion (Draper et al., 2004; Halliwell et al., 2020). Notably, chromosomes 1, 12, 17, 20 and X are particularly prone to full or partial gains in hPSCs (Baker et al., 2016; Draper et al., 2004; The International Stem Cell Initiative, 2011; Mayshar et al., 2010; Taapken et al., 2011). These aberrations often confer a growth advantage for the variant hPSCs, resulting in these cells eventually outcompeting the wildtype hPSC population in culture (Krivec et al., 2023; Nguyen et al., 2014; Stavish et al., 2023). However, at early cell passages it is often possible to find a mosaic population of variant and normal cells (Baker et al., 2007).

This genomic instability raises clear safety concerns when considering the use of hPSCs in clinical applications, as some of the recurrent CNVs are also present in cancer cells (Ben-David and Benvenisty, 2011; Halliwell et al., 2020). However, the impact of these genetic alterations on the utility of hPSCs for disease modelling has yet to be fully explored. While certain aberrations may not clearly affect the phenotype of undifferentiated hPSCs or their differentiation potential, they may have a more pronounced impact on differentiated cells, potentially altering cell function, physiology, and gene expression (Jo et al., 2020; Mehrjardi et al., 2020; Yang et al., 2010). This could compromise the value of hiPSCs as disease models for drug discovery if disease phenotypes are either obscured or even exacerbated. Notably the chromosome 1q region, which is recurrently gained, harbours several genes expressed in cardiac cells, including *TNNT2*, a gene encoding one of the major cardiac sarcomere proteins, and in which altered expression is known to result in cardiomyopathies (Vikhorev and Vikhoreva, 2018).

To determine whether a gain of the chromosome region 1q31.3-1q32.1, which includes *TNNT2*, would affect the cellular phenotype of a hiPSC line and its differentiated cardiomyocyte derivatives (hiPSC-CM), we carried out transcriptomic profiling and assessed the hiPSC-CMs for altered contraction kinetics. Our analysis revealed that hiPSC-CMs carrying a gain of this 1q region exhibited an enrichment of genes associated with the cardiac contraction machinery. Furthermore, these cells demonstrated a shorter contraction duration and irregular contractions when cultured in a multicellular three-dimensional model.

These findings underscore the critical need for genomic characterization and continuous monitoring of CNVs during extended culture of hPSCs. While these genetic changes may initially seem inconsequential to the culturing and differentiation of hPSCs, they can still impact the functionality of their differentiated derivatives, thereby influencing their suitability for disease modelling and drug response studies.

## RESULTS

### Human iPSCs with a gain of chromosome region 1q31.3 do not appear malignant

Continuous karyotype monitoring by SNP-based array on a control hiPSC line derived from a healthy individual (LUMC0020iCTRL-06; (Zhang et al., 2014)) detected a mosaic population with an ∼8MB duplication within chromosome region 1q31.3-1q32.1 (Figures 1A and S1A) that resulted in the duplication of 107 genes. Interphase FISH analysis revealed this gain to be an interstitial duplication, while notably G-band karyotyping failed to detect the duplication (Figures S1B and S1C). Digital droplet (dd)PCR confirmed the absence of the duplication prior to reprogramming, and that hiPSCs with the small chromosomal region gain (dup-1q31.3) gradually outcompeted the karyotypically normal wild-type hiPSC (WT-1q31.3) population (Figure S1D). Approximately 30% of the culture consisted of the dup-1q31.3 hiPSCs at an early passage, with this rising to 48% after 3 further passages, and finally establishing as a homogenous culture of 100% dup-1q31.3 cells within 17 passages.

**Figure 1.**
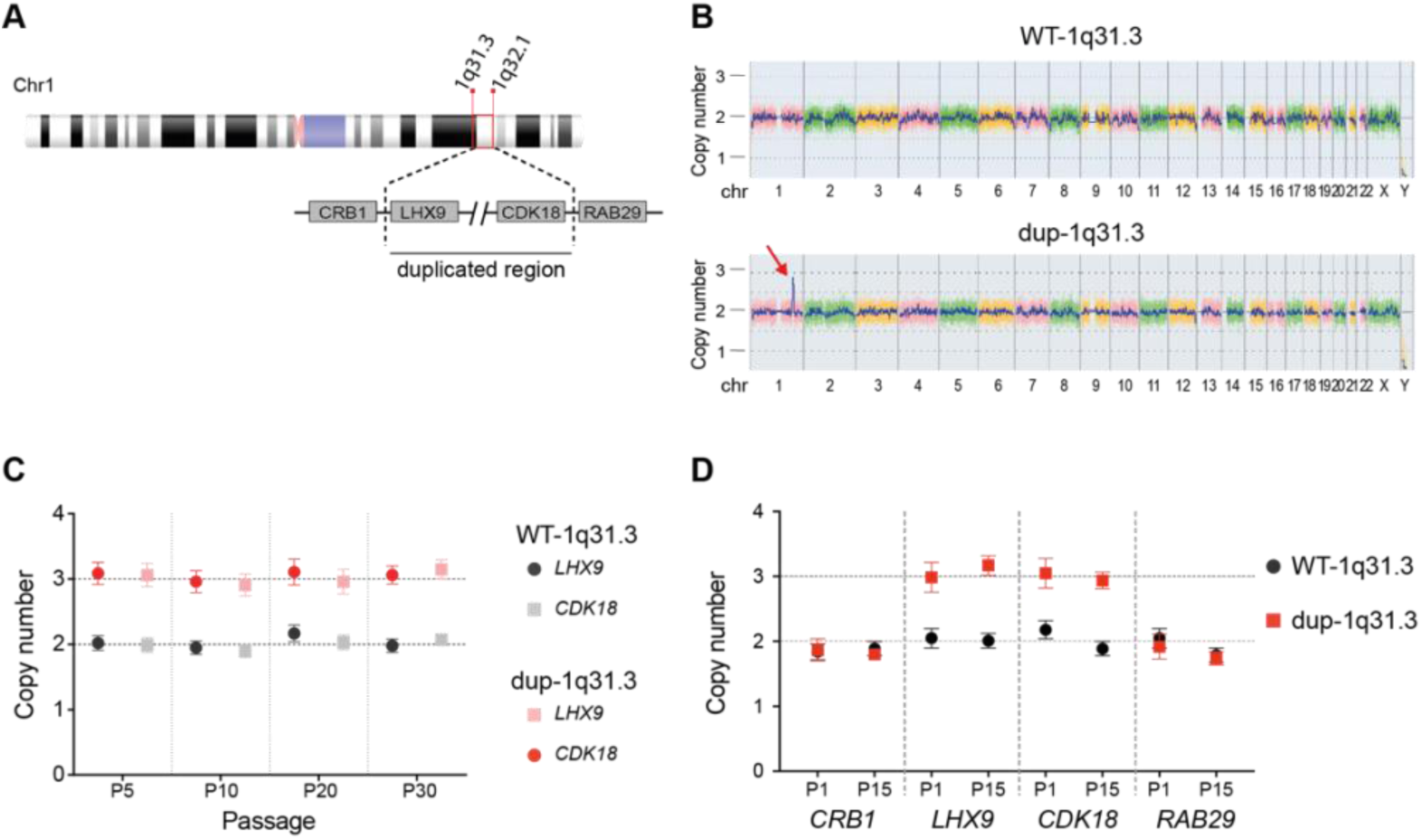
hiPSCs with a gain of 1q31.3-32.1 maintain a stable karyotype with prolonged culture. (**A**) Chromatid schematic highlighting the genes bordering the 7992kb duplicated region within 1q. (**B**) SNP-based microarray indicating the amplicon gain in the dup-1q31.1 hiPSCs (red arrow). **(C and D)** ddPCR-CNV assays for the WT-1q31.3 and dup-1q31.3 hiPSC lines indicating no further changes in number **(C)** or size of 1q31.3-32.1 region gained **(D)**. The copy number of genes at the 5’ and 3’ ends of the region duplicated (*LHX9* and *CDK18*), and the corresponding 5’and 3’ genes adjacent to the duplicated region (*CRB1* and *RAB21*) were assessed. P; passage number of the hiPSCs.

Following single cell dilution, clones with and without the 1q31.3-1q32.1 gain were isolated, as confirmed by SNP array analysis and ddPCR (Figures 1B and S1E). To determine if the subclones would maintain their respective genotypes with successive passaging, we assessed the hiPSCs for the chromosomal region gain by ddPCR every 5-10 passages. We confirmed genomic stability for the region in both cases for up to 30 passages. Moreover, the size of the gained amplicon and copy number remained constant during culture maintenance (Figures 1C and 1D).

To evaluate the potential malignancy of the 1q31.3-1q32.1 gain, we conducted an *in vivo* assay to determine if the hiPSCs would produce teratocarcinomas in mice (Figure 2A). Both hiPSC lines expressed the pluripotency markers OCT-4, SOX2 and SSEA-4 (Figure 2B) before subcutaneous transplantation into mice. Tumors were isolated for histopathological analysis between 4 and 12 weeks post-transplantation. Both lines yielded xenografts composed entirely of differentiated derivatives from the three embryonic germ layers without an undifferentiated component (Figure 2C), indicating that teratomas and not teratocarcinomas formed. Although the teratomas from dup-1q31.3 hiPSCs were harvested earlier due to reaching the maximum permitted size faster, there was no significant difference in average teratoma weight between the hiPSC lines (2125 ± 572mg and 3075 ± 790mg; WT-1q31.3 and dup-1q31.3, respectively), and no notable difference in the daily weight gain of the mice was observed (Figure 2D).

**Figure 2.**
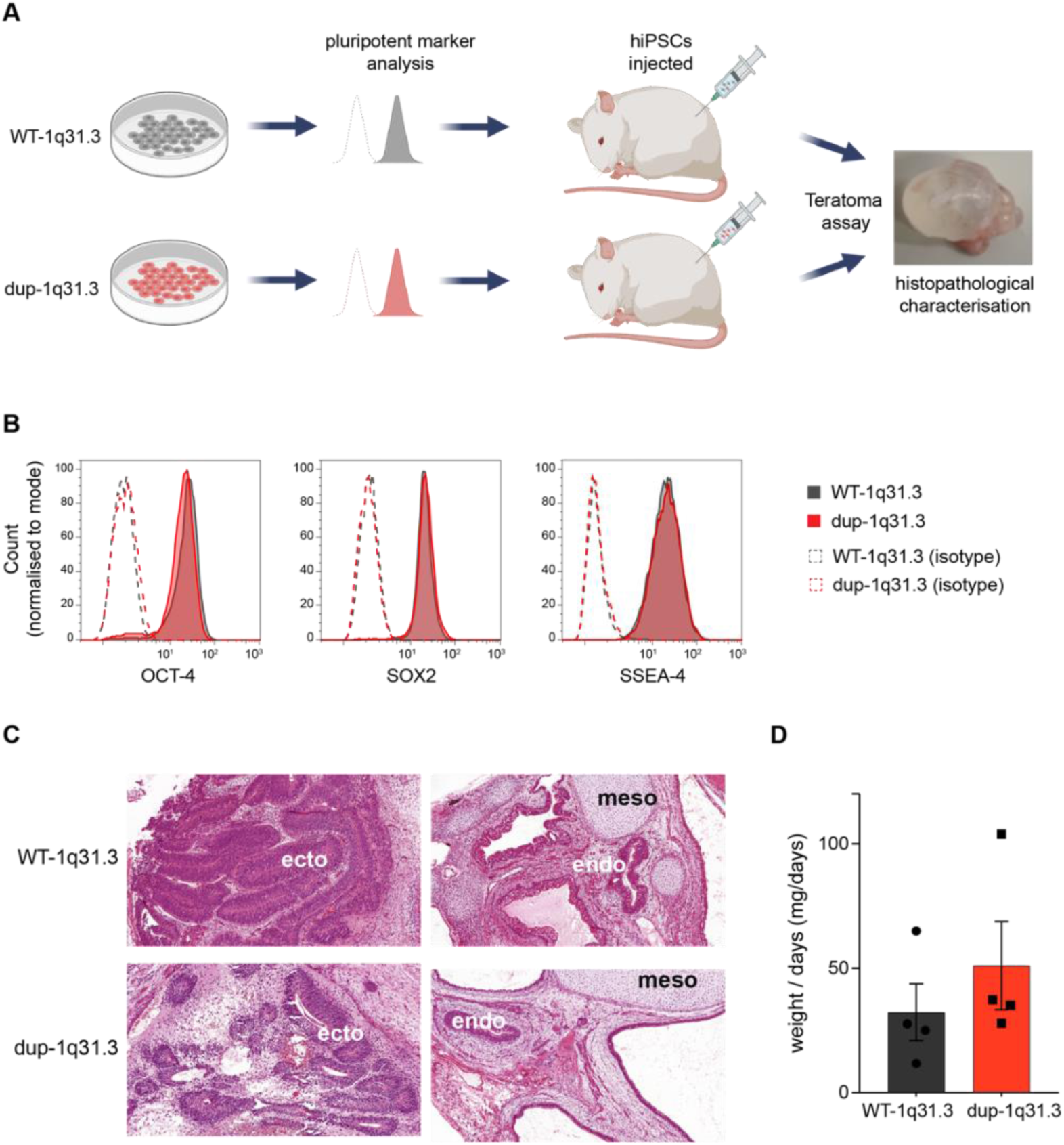
Histopathological examination indicates no differences in xenografts derived from WT-1q31.3 and dup-1q31.3 hiPSC lines. **(A)** Schematic representation of the workflow. WT-1q31.3 and dup-1q31.3 hiPSC were injected subcutaneously into immunodeficient mice (teratoma assay). Cell lines were evaluated for pluripotency markers before injection. At the endpoint, xenografts were collected for histological characterization. **(B)** Flow cytometry analysis of the pluripotency-associated markers OCT-4, SOX2 and SSEA-4 in the indicated hiPSC lines. Dotted lines indicate hiPSCs labelled with corresponding isotype control. **(C)** Representative hematoxylin and eosin-stained xenografts show differentiated tissue areas representing derivatives of mesoderm (meso), ectoderm (ecto) and endoderm (endo). **(D)** Bar graph indicating no difference in the average daily weight gain in the teratomas derived from the indicated hiPSC lines (n=4; P=0.23, Kolmogorov–Smirnov test).

### dup-1q31.3 hiPSC-CMs upregulate genes associated with contractility and cause a hypercontractile response in cardiac microtissues

We next explored the expression patterns of the 107 genes present in the gained 1q region. Filtering the duplicated region identified 46 genes (43% of the total CNVs) as being expressed in the human heart. Among these were genes involved in sarcomere organization (*TNNT2* and *LMOD1*). *TNNT2*, specifically, encodes cardiac troponin T (cTnT), a protein pivotal in regulating muscle contraction and an early cardiomyocyte marker during hPSC differentiation (van den Berg et al., 2015). We therefore investigated whether an additional copy of this gene affected the differentiation of the dup-1q31.3 hiPSCs to cardiomyocytes.

Flow cytometric analysis indicated comparable efficiency in cardiomyocyte differentiation between the lines, with approximately 77% of the differentiated cells expressing cTnT (Figure 3A). Moreover, there was no difference in the expression of MLC2v (∼40%), indicating that proportion of ventricular-like hiPSC-CMs was also similar between the two lines. However, RT-qPCR revealed an approximately twofold increase in *TNNT2* expression in dup-1q31.3 hiPSC-CMs compared to WT-1q31.3 hiPSC-CMs (P<0.001) (Figure 3B, left). Additionally, the mean fluorescence intensity (MFI) of cTnT confirmed significantly higher protein expression in the dup1q31.3 hiPSC-CMs (14250 ± 1722 vs 9844 ± 1340, respectively; P<0.02) (Figure 3B, right).

**Figure 3.**
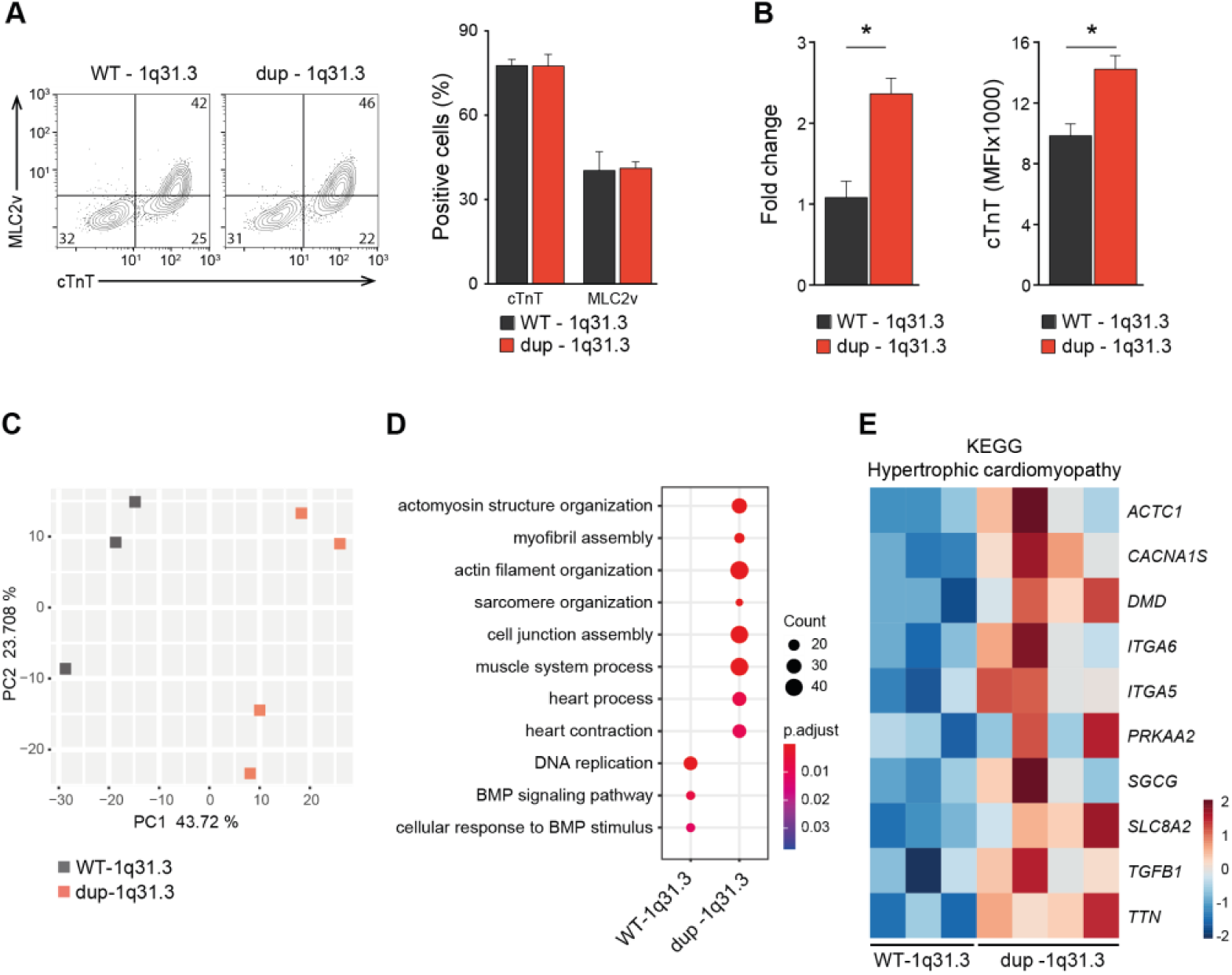
Gain of 1q31.3-32.1 upregulates expression of genes associated with contraction in hiPSC-CMs. **(A)** Representative flow cytometry plots (*left*) and overall cardiac differentiation efficiency (cTnT^+^) of WT-1q31.3 and dup1q31.3 hiPSCs (n=4) including average percentage that were ventricular-like (MLC2v^+^) cardiomyocytes (*right*). Values inside the dot plots are the percentage of cells within the gated region. **(B)** Expression level analysis of *TNNT2* by RT-qPCR (*left*) and cTnT by flow cytometry (*right*) in the WT-1q31.3 and dup-1q31.3 hiPSC-CMs (n=3; *P<0.05, unpaired t test). **(C)** PC analysis of RNA-seq replicate samples for WT-1q31.3 (n=3) and dup-1q31.3 (n=4) hiPSC-CMs. **(D)** GO terms enriched within DEGs between the WT-1q31.3 and dup-1q31.3 hiPSC-CMs. Colours represent the P_adjust_ value and dot size the number of genes mapped to the GO term (P_adjust_ < 0.05). **(E)** Heatmap of selected genes belonging to the hypertrophic cardiomyopathy KEGG pathway that are overall upregulated in dup-1q31.3 hiPSC-CMs.

To further understand the impact of the 1q region gain on hiPSC-CM phenotype, RNA-seq analysis was performed. Principal component (PC) analysis revealed distinctive differences between CMs derived from dup-1q31.3 hiPSCs and those from WT-1q31.3 hiPSCs (Figure 3C). Hierarchical clustering identified 1287 differentially expressed genes (DEGs) between the two lines, with 767 upregulated and 520 downregulated genes in the dup-1q31.3 hiPSC-CMs (Figures S2A and S2B, Table S2). While analysis of the RNA-seq data indicated that *TNNT2* was more highly expressed in the dup-1q31.3 hiPSC-CMs, it did not reach significance (RPKM WT-1q31.3 hiPSC-CM: 385 ± 42; RPKM dup-1q31.3 hiPSC-CM: 571 ± 71; P=0.057), most likely due to variability between the biological replicates (Figures 3C, S2C and S2D).

Gene ontology (GO) analysis highlighted that upregulated DEGs in dup-1q31.3 hiPSC-CMs were enriched in genes linked to muscle contraction machinery, while downregulated DEGs were associated with BMP signaling and DNA proliferation (Figure 3D). The upregulation of key genes involved in muscle cell differentiation, heart contraction and sarcomere organization indicates an augmented excitation-contraction coupling cascade (Figure S2E). Additionally, pathway annotation by Kyoto Encyclopedia of Genes and Genomes (KEGG) analysis revealed that genes associated with hypertrophic cardiomyopathy (HCM) were significantly upregulated in dup-1q31.3 hiPSC-CMs (Figure 3E).

We next investigated whether the altered expression of cTnT and other genes linked to sarcomere organization and cardiac contraction affected the contraction-relaxation kinetics in the dup-1q31.3 hiPSC-CMs. Representative contraction recordings of hiPSC-CM monolayers are depicted in Figure 4A. Analysis of the contractility kinetics using MUSCLEMOTION revealed no significant differences in contraction duration (ContD), time to peak, relaxation or amplitude between WT-1q31.3 and dup-1q31.3 hiPSC-CMs. (Figure 4B).

**Figure 4.**
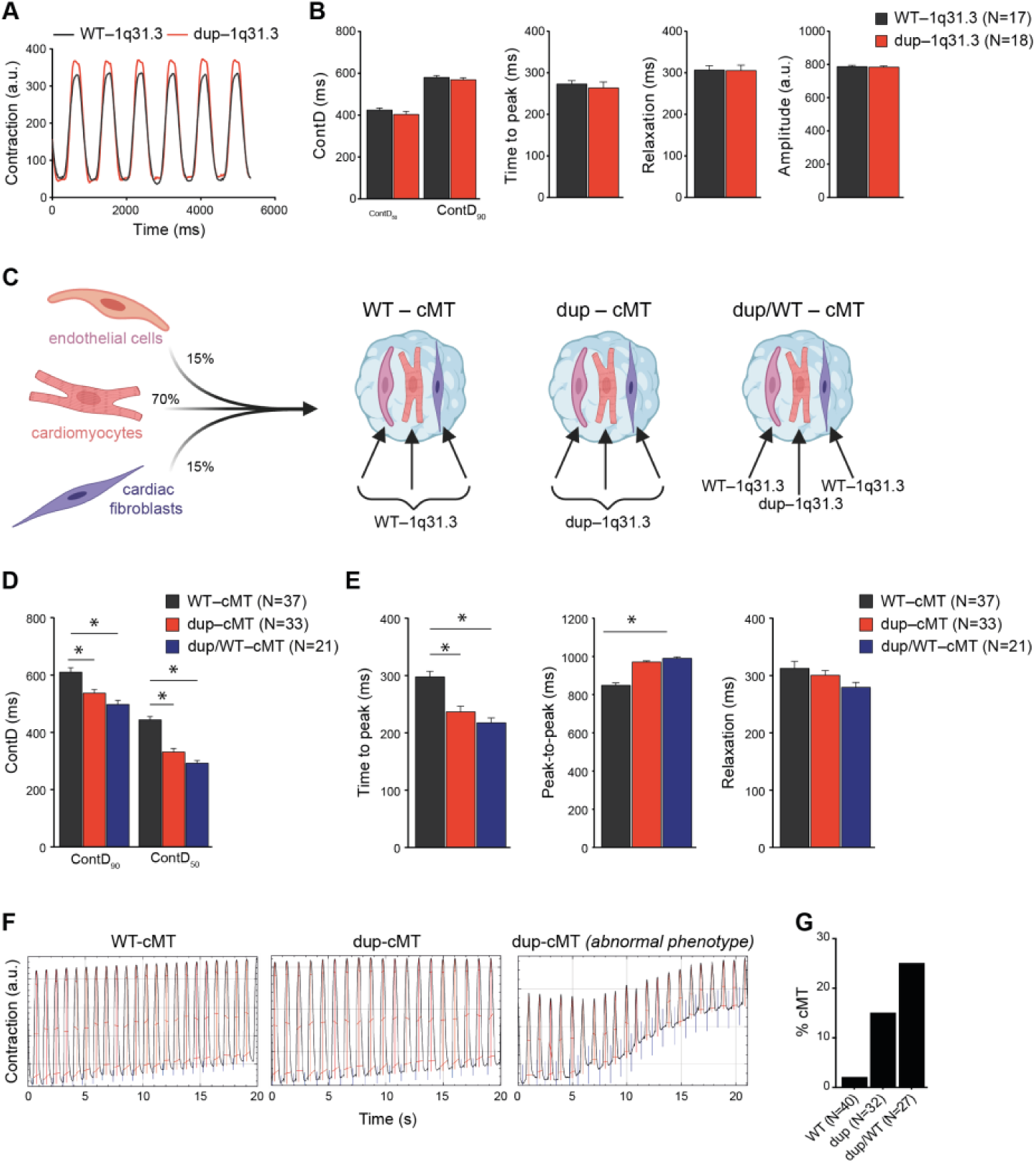
dup–1q31.3 hiPSC-CMs exhibit hypercontractility in cardiac microtissues (cMTs). **(A)** Representative contraction traces from paced (1 Hz) WT-1q31.3 (black) and dup-1q31.3 (red) hiPSC-CMs when cultured as monolayers. **(B)** Quantitative analysis of contractile properties for the indicated hiPSC-CM lines when cultured as monolayers. Parameters include contraction duration at 50% and 90% (ContD_50_ and ContD_90_), contraction (time-to-peak) and relaxation time, and amplitude (n=4 differentiations). Values (N) refer to the number of wells analyzed. **(C)** Schematic of the three different types of cMT created, with the corresponding percentage and genotype of each cell type. **(D and E)** Quantitative analysis of contractile properties for the indicated cMTs including average ContD_50_ and ContD_90_ **(D),** time-to-peak, peak-to-peak and relaxation **(E)** values (n=3 differentiations; *P<0.05, unpaired t test). Values (N) refer to the number of cMTs analyzed. **(F)** Representative contraction traces from WT- and dup-cMTs. **(G)** Bar graph summarizing the percentage of cMT from each group displaying an abnormal contraction profile (P=0.0120, Fisher’s exact test). Values (N) refer to the number of cMTs analyzed.

The absence of discernible differences in contractile responses in the dup-1q31.3 hiPSC-CMs despite the altered expression of genes related to cardiac contraction might be attributed to hiPSC-CM immaturity. To determine whether maturation in 3D cultures containing other cardiac cell types would reveal differences in contraction, we generated cardiac microtissues (cMTs) by co-culturing the hiPSC-CMs with CNV-matched hiPSC-derived endothelial cells (hiPSC-ECs) and cardiac fibroblasts (hiPSC-CFs) (Figure 4C). Additionally, we created cMTs by aggregating the dup1q31.3 hiPSC-CMs with WT-1q31.3 hiPSC-ECs and -CFs (dup/WT cMT). Subsequent MUSCLEMOTION analysis of paced cMTs revealed a significant decrease in ContD in dup-1q31.3 cMT compared to WT-1q31.3 cMT (Figure 4D). This difference was also evident in dup/WT cMT, suggesting that the hiPSC-CMs were responsible for the hypercontractile phenotype. Examination of additional parameters indicated that the decrease in ContD was due to a faster time to peak, while relaxation remained consistent across all three types of cMTs (Figure 4E).

Blinded examination of the contraction traces revealed irregular contractions in some of the dup- and dup/WT-cMTs (Figure 4F). This irregular phenotype, characterized by progressive shortening of ContD and amplitude during recording, was more frequently observed in cMTs containing dup-1q31.3 hiPSC-CMs (Figure 4G). Collectively, these findings suggest that hiPSC-CMs overexpressing TNNT2 exhibit impaired contraction, potentially contributing to the pathological phenotype observed in the cMTs.

## DISCUSSION

The influence of karyotypic irregularities on the utility of hPSCs in various biomedical applications is a growing area of focus. These acquired genetic alterations which can emerge rapidly and may not be immediately evident, not only raise safety concerns for the use of hPSC-based therapies in regenerative medicine but may also impact broader applications such as disease modelling and drug discovery. Detecting these variants is pivotal for ensuring the quality of hPSCs.

In this study, we identified a hiPSC line that at low passage consisted of a mosaic population comprised of both wild type cells and cells with a gain of the chromosome region 1q31.3-1q32.1. Significantly, this mosaicism eluded detection through G band karyotyping, highlighting the challenges of detecting relatively small size CNVs and the need to employ multiple approaches to confirm the integrity of karyotypes. Moreover, *de novo* CNVs may be present in the human somatic cells themselves prior to reprogramming (Abyzov et al., 2012; Jacobs et al., 2014), indicating a high degree of genetic variability that might still be underestimated in hPSCs.

Over extended passages, the cells harbouring an interstitial duplication in amplicon 1q31.3 gained a selective growth advantage and eventually dominated the hiPSC culture, although it was possible to isolate and maintain the wild type hiPSCs prior to this occurring. Aberrations in chromosome 1q are recurrent in hPSCs, with the prevalence of this abnormality increasing in recent years and correlating with a shift from the culturing of hPSCs on mouse embryonic fibroblasts (Stavish et al., 2023). Recent studies have reported that the anti-apoptotic gene *MDM4,* located in the duplicated chromosomal region, as the likely driver for the competitive advantage of hPSCs containing a gain of chromosome 1q (Krivec et al., 2023; Stavish et al., 2023). *MDM4* suppresses the p53 response to cellular stress, thereby increasing the threshold for triggering apoptosis. Moreover, duplications of regions within 1q are detected in many cancers (Guichard et al., 2012; Schmidt et al., 2021), with MDM4 also proposed to contribute to the frequent occurrence of chromosome 1q amplifications in breast cancer (Girish et al., 2023).

Despite this, transplantation of the dup-1q31.3 hiPSCs into immunocompromised mice did not result in malignancy. While it could be expected that the dup-1q31.3 hiPSC line would be more likely to generate teratocarcinomas due to their enhanced proliferative potential, a recent International Stem Cell Initiative (ISCI) study established no clear correlation between teratocarcinoma formation and karyotypic abnormalities (The International Stem Cell Initiative, 2018).

Similarly, the ability of the dup-1q31.3 hiPSCs to differentiate into ventricular-like cardiomyocytes remained unaffected. This lack of effect on differentiation capacity is likely due to the differentiation protocol used where notably the concentration of CHIR99021, a Wnt signalling pathway activator, is lower compared to some other published differentiation protocols (Lian et al., 2012), and possibly because both a tankyrase inhibitor (XAV939) and a porcupine inhibitor (IWP-L6) subsequently are used to down-regulate Wnt signalling (Brink et al., 2020).

While cardiomyocyte differentiation of dup-1q31.3 hiPSCs was unaltered, the duplication of genes involved in muscle contraction led to increased expression of sarcomere genes (e.g. *ACTC1, TTN*) and *TGF-β1*, a mediator of cardiac hypertrophy. The duplication of *TNNT2* also resulted in both increased transcript and protein expression in the hiPSC-CMs. Although missense mutations in *TNNT2* are more frequently observed, there is a documented case of an HCM patient with an extra copy of *TNNT2* (Lopes et al., 2015), with features of the disease including impaired diastolic function and hypercontractility.

Although transcriptomic and proteomic changes were observed in hiPSC-CMs when cultured in 2D, these did not manifest a contractility phenotype. However, in a 3D culture system known to promote the maturation of hiPSC-CMs (Giacomelli et al., 2020), faster as well as irregular contractions were observed in cMTs containing dup-1q31.3 hiPSC-CMs. Previous studies have also observed different degrees of HCM-associated contractility abnormalities with sometimes conflicting results when the hiPSC-CM HCM models were analysed in either 2D or 3D (Guo et al., 2021). Compared to 2D systems, 3D models support the development of more mature and functional sarcomeres (Drakhlis and Zweigerdt, 2023), highlighting the importance of physiologically relevant *in vitro* model systems. The differences between the 2D and 3D culture models may also be attributed to the rigidity of the substrate affecting force generation.

In summary, this study underscores the importance of careful monitoring for chromosome aberrations in hPSC lines, not only in the context of cell therapy but also potentially in research settings like disease modelling or drug screening. Although these abnormalities might not manifest observable changes in undifferentiated cell populations, they could significantly confound disease phenotype readouts, Investigating the implications of such abnormalities on other differentiated hPSC derivatives also warrants further investigation.

## EXPERIMENTAL PROCEDURES

### Human induced pluripotent stem cell (hiPSC) lines

The hiPSC lines used in this study the hiPSC line, LUMC0020iCTRL (female, (Zhang et al., 2014), RRID: CVCL_ZA25), and subclones of this line (WT-1q31.3 and dup-1q31.3). The hiPSCs were maintained in StemFlex™ Medium (Gibco) on Laminin-521 (Biolamina)-coated plates, as previously described (Brandão et al., 2020). While the UMC0020iCTRL and dup-1q31.3 hiPSCs appeared karyotypically normal by G-banding, a duplication of 1q31.3-1q32.1 on chromosome 1 was detected in some or all the cells by SNP array analysis or by ddPCR.

### Differentiation of hiPSCs and generation of cardiac microtissues (cMTs)

The hiPSC-derived cardiomyocytes (hiPSC-CMs), endothelial cells (hiPSC-ECs) and cardiac fibroblasts (hiPSC-CFs) were differentiated and cMTs generated as previously described (Campostrini et al., 2021). Analysis of cMTs was performed between 3 and 4 weeks after formation. Three independent cardiomyocyte differentiations were used per line.

### Flow cytometric analysis

hiPSCs or hiPSC-CMs were prepared for flow cytometric analysis as previously described (Brandão et al., 2020). The list of antibodies used is provided in Supplemental Information.

### Teratoma assay

The teratoma assay was performed as described previously (Salvatori et al., 2018). Between 4-12 weeks after injection, the tumours were removed, weighed, and subjected to histopathological analysis. Animal experiments were approved by the Dutch Central Commission for Animal Experimentation (Centrale Commissie voor Dierproeven).

### Gene expression analysis

For both RT-qPCR and RNAseq analysis a minimum of 3 biological replicates for each sample were analysed.

### Contraction analysis of hiPSC-CMs

Both hiPSC-CM monolayers and cMTs were analysed using the MUSCLEMOTION ImageJ macro (ImageJ) (Sala et al., 2018). At least three independent differentiations per line were used.

## Author Contributions

Conceptualization, K.O.B. and R.P.D.; Methodology, K.O.B., V.M., D.S., and R.P.D.; Investigation, K.O.B., V.M., L.Y., and D.S.; Formal Analysis, K.O.B., X.C., L.S., D.S. and R.P.D.; Resources: M.P.H.M; Supervision, M.B., C.L.M. and R.P.D.; Writing – Original Draft, K.O.B. and R.P.D.; Writing – Review & Editing, all authors; Funding Acquisition, M.B., D.S., C.L.M. and R.P.D.

## Acknowledgements

This work was supported by a Starting Grant (STEMCARDIORISK) from the European Research Council (ERC) under the European Union’s Horizon 2020 Research and Innovation Programme (H2020 European Research Council; grant agreement no. 638030), a VIDI fellowship from the Netherlands Organisation for Scientific Research (Nederlandse Organisatie voor Wetenschappelijk Onderzoek NWO; ILLUMINATE; no. 91715303), and by a Novo Nordisk Foundation grant (NNF21CC0073729; reNEW). Some panels within figures were created with BioRender.com.

---

**Figure S1.**
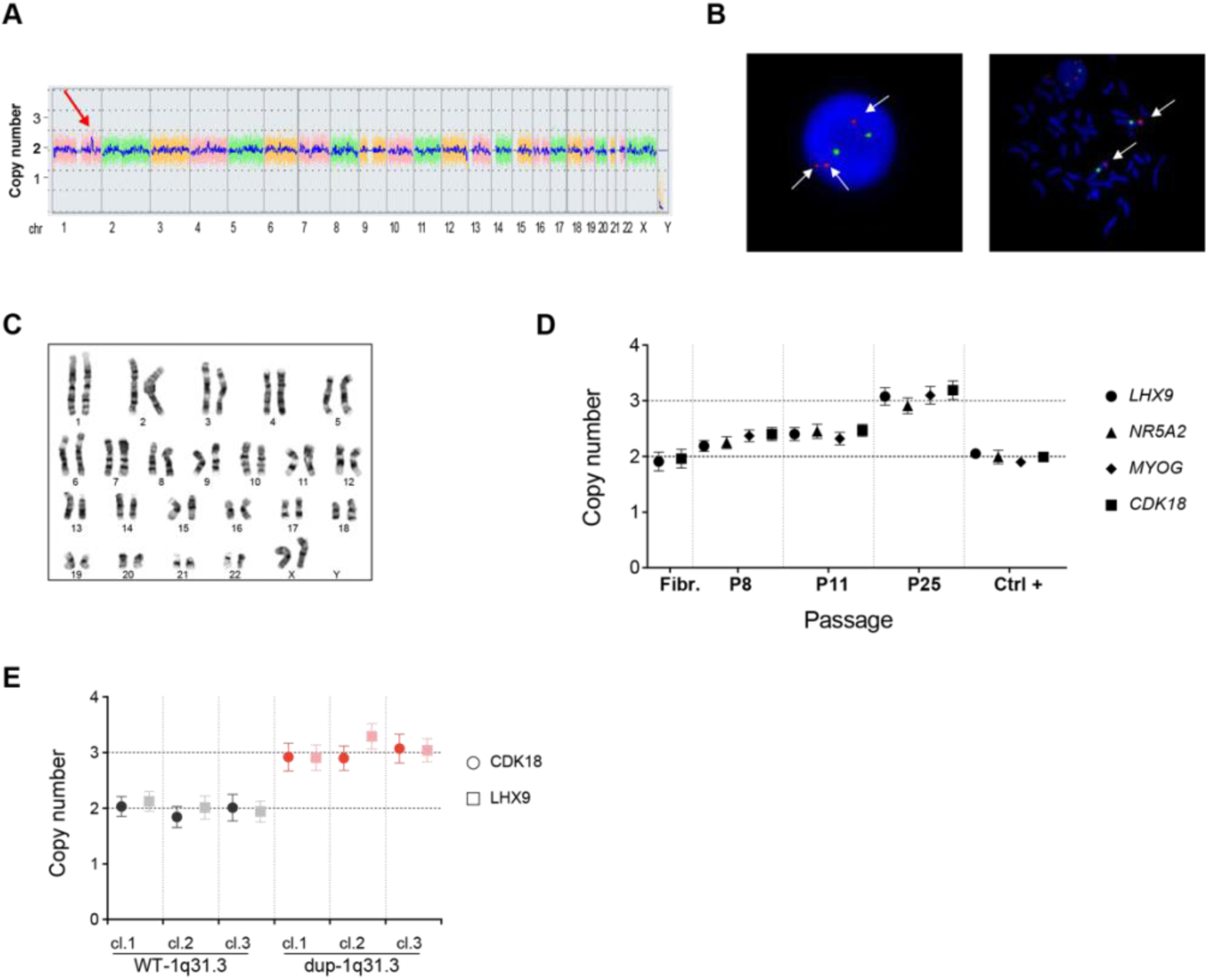
Identification of a mosaic population of hiPSCs with a gain of 1q31.3-32.1 and subsequent subcloning. Related to Figure 1. (**A**) SNP-based microarray indicating some cells in a population of the hiPSC line LUMC0020iCTRL show a copy gain of 1q31.3-32.1, as indicated by the red arrow. (**B**) Dual-colour FISH analysis with MDM4 probe at 1q32 (red) and satellite DNA probe at 1q12 (green). DAPI was used as the nuclear counterstain. (**C**) G-banding karyogram is unable to detect a copy gain of 1q31.3-32.1. (**D**) ddPCR-CNV assay demonstrating the gradual increase of hiPSCs within the population showing a CNV gain of 1q31.3-32.1 with continued maintenance in culture. The copy number of 4 genes located in the 1q31.3-32.1 region were assessed. The fibroblasts (Fibr.) used for reprogramming to hiPSCs, as well as an independent hiPSC lines (LUMC0099iCTRL; Ctrl+) did not appear to show CNV for this region. (**E**) ddPCR-CNV assay identifying subclones of the mosaic LUMC0020iCTRL hiPSCs containing 2 (WT-1q31.3) or 3 (dup-1q31.3) of the 1q31.3-32.1 region. *LHX9* and *CDK18* are the respective 5’ and 3’ genes at the border of the duplicated region.

**Figure S2.**
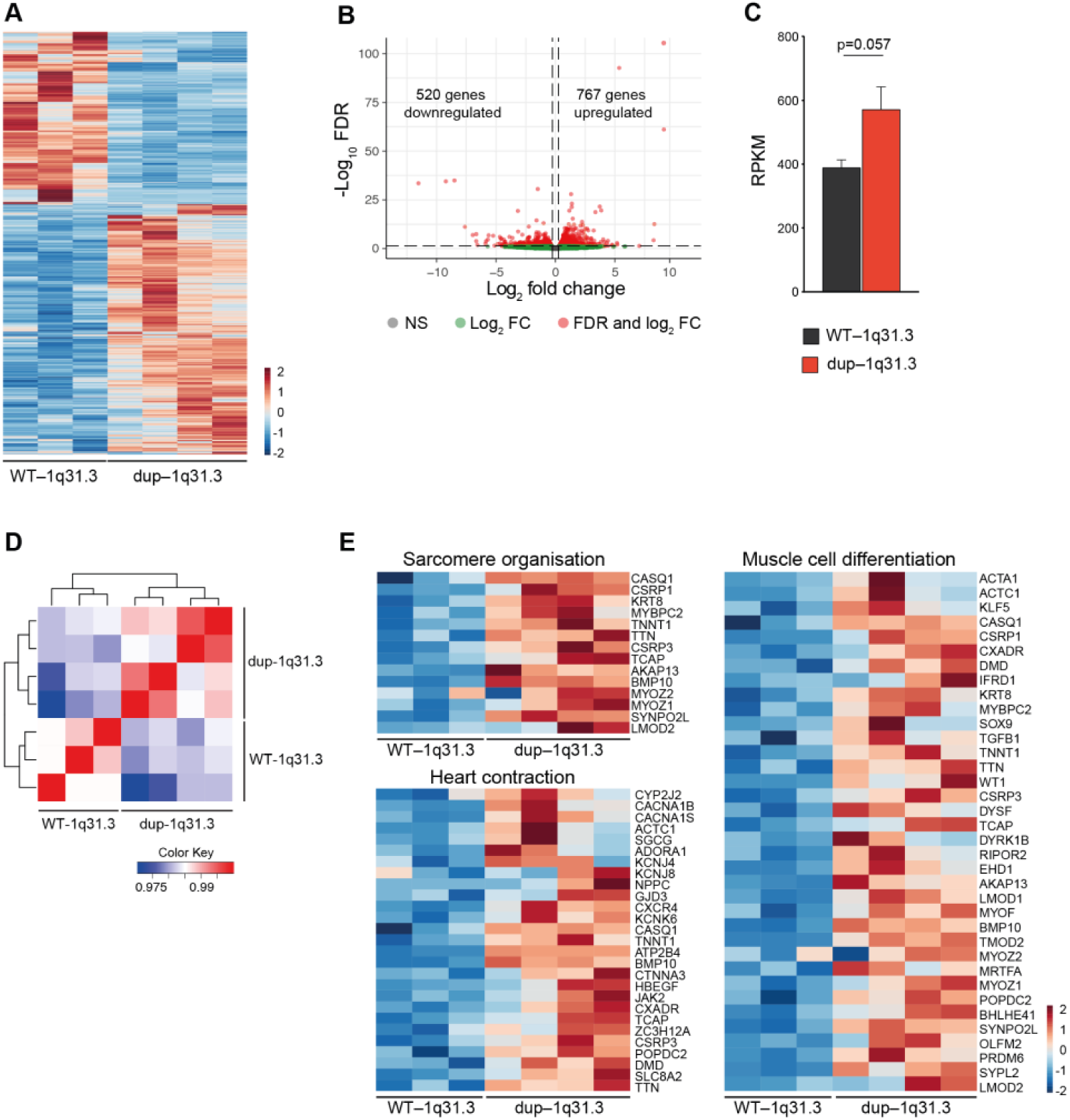
Gain of 1q31.3-32.1 upregulates expression of genes associated with contraction in hiPSC-CMs. Related to Figure 3. **(A)** Heatmap showing differentially expressed genes (DEGs) identified in the replicate WT-1q31.3 and dup-1q31.3 hiPSC-CM samples (1287 genes, FDR < 0.05). **(B)** Volcano plot showing sorted log_2_ fold change (FC) for dup-1q31.3 hiPSC-CMs. Log_2_ FC > 0 and log_2_ FC < 0 respectively indicate upregulated and downregulated genes in dup-1q31.3 hiPSC-CMs compared to WT-1q31.3 hiPSC-CMs. **(C)** Average RPKM value for *TNNT2* in WT-1q31.3 and dup-1q31.3 hiPSC-CMs from RNA-seq analysis. **(D)** Spearman’s correlation heatmap of WT-1q31.3 and dup-1q31.3 hiPSC-CMs based on RNA-seq data. **(E)** Heatmap of genes in 3 selected GOs (from Figure 3D) related to contraction enriched in the upregulated DEGs of dup-1q31.3 hiPSC-CMs.

